# The effect of selective logging on microclimates, arthropod abundance and the foraging behaviour of Eastern Himalayan birds

**DOI:** 10.1101/2022.08.24.504948

**Authors:** Kanika Aggarwal, Ritobroto Chanda, Shambu Rai, Mangal Rai, D.K. Pradhan, Binod Munda, Bharat Tamang, Aman Biswakarma, Umesh Srinivasan

**Affiliations:** Institute of Environment and Sustainable Development, Banaras Hindu University, Varanasi, Uttar Pradesh 221005, India; Centre for Ecological Sciences, Indian Institute of Science, Bangalore 560 012, India

**Keywords:** arthropod density, arthropod diversity, foraging success, forest degradation, gleaner, land-use change, microclimate, montane birds, sallier, time spent foraging, understorey insectivores

## Abstract

1. Selective logging—the practice of removing a subset of commercially important trees from a forest—is a globally pervasive form of forest degradation. Selective logging alters both the structure and function of forests and the composition of ecological communities.

2. Tropical insectivorous birds are highly vulnerable to microhabitat alterations in logged forest. Such altered microhabitats might affect the foraging of forest birds by altering (a) resource availability, and (b) foraging behaviour.

3. We investigated the effect of selective logging on microclimates, prey availability, foraging behaviour and the foraging success of eastern Himalayan birds in the breeding season.

4. Selective logging alters temperature-humidity microclimates and the composition of arthropod communities, both of which are likely to then collectively alter foraging behaviour by birds. We show that birds spent a lower proportion of their time foraging in primary compared with logged forest. Further, selective logging interacts with species traits such as body mass, preferred foraging stratum (understorey, midstorey or canopy) and foraging manoeuvre to influence foraging success. Gleaners generally foraged more successfully in primary forest and salliers in logged forest, although these patterns were modified by body mass and foraging stratum.

5. Synthesis and applications: Our study shows how altered microclimates in anthropogenically modified habitats can influence resource availability and have downstream impacts on the behaviour of species at higher trophic levels.

## INTRODUCTION

Forests cover 31% of our planet’s land and harbour about 80% of terrestrial plant and animal biodiversity (FAO & UN GFGR, 2021). Amongst forests worldwide, tropical forests are especially biodiverse, and even within the tropics, forested habitats in tropical mountains harbour a disproportionately high share of biodiversity (Elsen et al., 2018). However, tropical montane species are increasingly under threat from habitat loss and degradation as well as climate change (Pimm, 2008; Freeman et al., 2021), and both these drivers of biodiversity loss can interact with each other (with habitat degradation complicating the abiotic impacts of climate change) to affect the behaviour and fitness of species (Srinivasan & Wilcove, 2021).

Of the various ways in which tropical forests are lost or degraded, selective logging—the practice of removing a subset of commercially important trees from a forest—is especially pervasive. A total of 20% of tropical forest was logged in just a five-year period and selective logging continues to be the most widespread form of tropical forest degradation (Edwards & Laurance, 2013). Logging creates gaps in the forest canopy and allows direct sunlight to reach the lower levels (understorey and midstorey) within the forest (Senior et al., 2018). This often leads to warmer and drier conditions in logged than in primary forest and changes in the availability of microhabitats that might be important for a variety of species (Senior et al., 2017).

Tropical rainforests host roughly six million invertebrate species (Hamilton et al., 2010) and over 18,000 species per hectare (Basset et al., 2012; Ewers et al., 2015), and the structural and environmental changes that arise from selective logging can impact patterns of invertebrate diversity and abundance. Logging has been shown to reduce the abundance of key invertebrate decomposers (such as leaf-litter beetles, and termites) to two-thirds of their abundance in primary forest (Ewers et al., 2015). Altered microclimatic conditions in logged forest, such as higher temperatures and lower humidity most likely explain reductions in the abundance or biomass of invertebrates in a logged forest (Ewers et al., 2015), as soft-bodied invertebrates are particularly sensitive to desiccation (Cornelius et al., 2010). Logging, therefore, creates important changes to arthropod communities (Holloway et al., 1992; Hill et al., 1995; Hill, 1999; Willott et al., 2000; Vasconcelos et al., 2000; Davis, 2000; Basset et al., 2001). In general, logged forests possess lower arthropod species richness, and a few species dominate arthropod communities that possess high tolerance potency towards logging-induced altered microclimates (Basset et al., 2001).

Selective logging also alters the structural and functional composition of bird communities (Burivalova et al., 2015). Amongst tropical birds, insectivorous species (whose diets are dominated by arthropods) are especially sensitive to land-use change (Bregman et al., 2014; Srinivasan et al., 2015; Powell et al,. 2015). Across the tropics, studies have repeatedly found that terrestrial insectivorous are the most vulnerable to changes in forest structure, and are often the first dietary guild to disappear from disturbed forest (Stratford & Stouffer, 1999; Canaday & Rivadeneyra, 2001; Peh et al., 2005; Pavlacky et al., 2015; Rutt et al., 2019; Stouffer et al., 2021) and the last to return after forests regenerate (Powell et al., 2013, 2015).

Microclimatic changes due to selective logging which include high sunlight penetration and high temperature are induced by edge effect and might make logged forest patches unsuitable for physiologically detrimental to birds (Stratford & Robinson, 2005). Altered microhabitats could also affect foraging by insectivorous birds indirectly by altering resource availability and diversity (Powell et al., 2015).

We asked how selective logging affected (a) microclimates, (b) arthropod availability for insectivorous birds, and (c) the potential impacts these might have on the foraging behaviour of Eastern Himalayan insectivorous birds. Globally, the Himalayas are amongst the most biodiversity-rich terrestrial regions (Grenyer et al., 2006) and by 2100, they are at risk of losing approximately half of their forest cover to land selective logging and other forms of land-use change (Pandit et al., 2007). We hypothesised that higher temperature and lower humidity with logging would alter the abundance and diversity of arthropod prey (i.e., resources) for insectivorous birds in logged forest. Further, we hypothesized that increased temperatures in a logged forest would also affect the time spent foraging by birds, as higher temperatures might prevent birds from foraging actively.

We predicted that:

(a) In line with prior work, logged forest would be warmer and drier than primary forest.
(b) Altered microhabitats would be associated with changes in the densities of various kinds of arthropods.
(c) As a direct impact of altered microclimates, birds would spend less time foraging in logged forest versus primary forest because of potential thermal stresses associated with activity in warmer environments.
(d) As a potential indirect impact of altered microclimates mediated via changes in arthropod abundance, foraging success of birds would be altered depending on whether logging increases or decreases the abundances of their particular arthropod prey types.
(e) Finally, we expected that species traits such as body mass and foraging stratum would affect foraging success, driven by the differences in resource requirements of birds of different sizes and by the availability of arthropod prey at understorey, midstorey and canopy level.

## METHODS

### Study Area

We conducted fieldwork in Eaglenest Wildlife Sanctuary (EWS), West Kameng district, Arunachal Pradesh, India (27.07°N; 92.40°E; Fig. 1). The study area was located in the tropical montane broadleaved forest at 2,000m above sea level. Within this habitat, the canopy is dominated by tree species from the genera *Quercus, Betula, Acer, Michelia* and *Alnus*, with bamboo (*Chimonobambusa sp*.) and ferns in the understorey. Parts of the forest at this location were selectively logged until 2002, after which logging was stopped. Tree densities in the primary forest plots (> 30cm DBH; 168 to 192 trees ha^-1^) are roughly two to two-and-a-half that in logged forest plots (76 to 109 trees ha^-1^). The fragmentation of the canopy has resulted in the increase of bamboo in the understorey (primary forest, bamboo stem density = 0.37m^-2^ ± 0.07SE; logged forest, bamboo stem density = 0.94m^-2^ ± 0.22SE), which has since hampered the propagation of forest tree saplings in the understorey, such that vegetation structure in these sampling plots has not changed since 2011 (Srinivasan and Wilcove, 2021). We sampled primary and logged forest in four plots, two each primary and logged forest (Fig. 1). The total area sampled in the primary and logged forest were 6 ha and 6.5 ha respectively.

**Figure 1.**
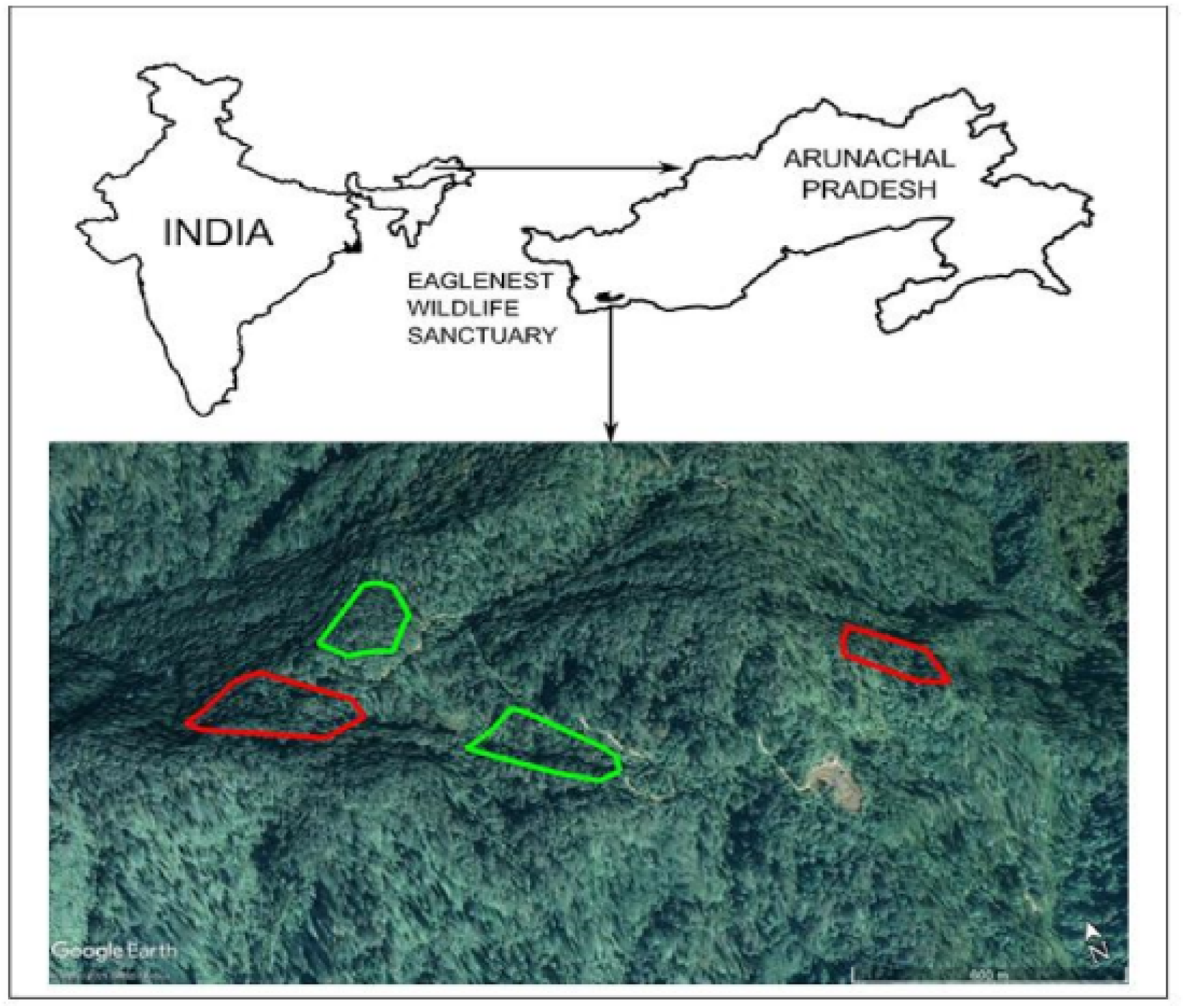
Map of the study area in Eaglenest Wildlife Sanctuary, Arunachal Pradesh, India. Sampling plots in primary forest are outlined in green and plots in logged forest are outlined in red.

### Field Methods

#### Measuring temperature and humidity

Within each of the primary and logged sampling plots, we placed 20 automated temperature and humidity loggers (Maxim Integrated iButton Hygrochron DS1923-F5#) in a grid-based pattern, with ∼50m between successive loggers. At each location, each logger recorded temperature and relative humidity every half hour during the day. We then downloaded the temperature and humidity data to compare these abiotic parameter values between primary and logged forest.

#### Arthropod Data collection

In each primary and logged forest plot, we placed 25-28 sampling stations, with neighbouring stations spaced roughly 40m apart. At each station, we sampled terrestrial, foliage and flying arthropods using, respectively, a single pitfall trap placed for 48 hours, 10 beats to a single branch taken at the location and a single sticky trap placed for 48 hours. Each pitfall trap was a small plastic container (5 cm diameter; 7cm in height) filled with detergent water and buried in the ground up to lip-level. For branch beating we stitched a white linen cloth in the shape of a funnel around a metal ring 1m in diameter at the larger end of the funnel and with a container attached to the thinner end of the funnel. At each station, we placed a randomly selected branch within this funnel, beat it ten times with a standard-sized stick and sprayed commercially available insecticide spray to momentarily paralyze the arthropods, which were collected in the container affixed to the narrow end of the funnel. The sticky trap for flying arthropods was a commercially available plastic sheet (150 × 200mm) with a blue background and glue on both sides (Chipku brand). We hung the sticky trap from a branch at the station 1.5m above the ground. We collected all arthropods from each sample and photographed them against a white background with a scale in the photograph. We then identified each arthropod to the order/class level.

#### Bird foraging behaviour

We collected data on bird foraging behaviour from 8 March to 30 April 2021 in two-time intervals each day (0700 to 1100hrs and 1400 to 1600hrs). We sampled such that primary and logged forest plots were equally sampled during the morning and afternoon. Once within a sampling plot, we actively searched for birds using either visual or acoustic cues. Upon seeing a bird, we observed it using binoculars (Zeiss Terra ED 8×42) for the time period when it was clearly visible. For each such observation, we noted (a) start time and end time (to calculate the duration of the observation in seconds), (b) species identity, (c) foraging height using a laser rangefinder (Hawke LRF400), (d) the number of foraging attempts made by the individual, and whether each attempt was a success or a failure, (e) the food item that was targeted during each attempt, (f) the substrate from which the food item was captured, and (g) the foraging manoeuvre used to attempt each instance of feeding. We ended each observation when the bird was no longer clearly visible. We classified a foraging attempt as a success if we observed the bird swallowing after making a foraging attempt. Along with foraging, birds were also observed involved in other non-foraging behaviours such as preening and calling.

Foraging manoeuvres were classified following (Robinson & Holmes, 1982) as hover (bird captures a stationary food item while it is itself in flight), glean (stationary bird capture stationary prey) or sally (flying prey is pursued and captured while the bird is in flight) was also recorded. Weather updates were recorded every half an hour in the form of presence or absence of sun, cloud, fog and rain during fieldwork. Data from the first five days of fieldwork were considered trial data and therefore excluded from the analyses.

### Analytical Methods

#### Temperature and relative humidity

We pooled all the temperature and relative humidity values in logged forest and primary forest patches separately. To compare temperatures between the two habitats, we selected temperature records between 11 AM and 12 PM (mid-day). We then used an ANOVA to estimate temperature differences between logged forest and primary forest. We also used an ANOVA to compare relative humidity values between logged forest and primary forest.

#### Arthropod numbers and selective logging

We calculated arthropod densities after combining the data from the pitfall trap, branch beat and sticky traps at each station. We used generalized linear models (GLMs) with categorical habitat types (primary or logged) to model the impact of logging on arthropods.

#### Arthropod community composition

We used non-metric multidimensional scaling (NMDS) ordination analysis to compare order-level arthropod community composition in primary and logged forest using the Morisita-Horn index in the R package *vegan* (Okansen et al., 2019; R Core Team 2021). For the NMDS analysis, each “site” was an arthropod sampling station, and for each station, we combined the arthropod data from the pitfall and sticky traps and branch beats. Therefore, aach point in the NMDS visualisation represents the composition of arthropods obtained from the three sampling methods at a single sampling station. We excluded arthropod orders represented by fewer than ten individuals collected from the analysis.

#### Time spent foraging

For individual species with at least ten observations (49 species), we calculated the proportion of time spent foraging (total time in seconds the species was observed actively foraging, divided by the total time the species was observed foraging and engaged in other activities such as preening) in primary and logged forest separately. This value theoretically ranges from 0 (no time spent foraging) to 1 (100% of time spent foraging). To test the factors that might affect time spent foraging, we ran a generalised linear model of the quasibinomial family (because the response variable was a ratio bound between 0 and 1) with the following formulation:

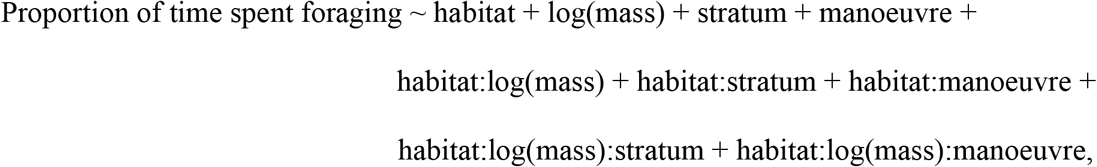

Where habitat was a factor variable with two levels (primary and logged), stratum was a factor variable with three levels (understorey, midstorey and canopy) and the manoeuvre was a factor variable with two levels (sally and non-sally; gleaners and hover-feeders were clubbed into a single category of birds feeding on prey in vegetation). Semicolons in the formulation represent interactions between predictor variables.

#### Foraging success

For each species with at least 10 observations in primary and logged forest separately, we first selected only the observations in which an individual was actively foraging. For each such observation, we calculated the number of successful and failed foraging attempts – this information was used as a response variable in a binomial generalised mixed model with the following formulation:

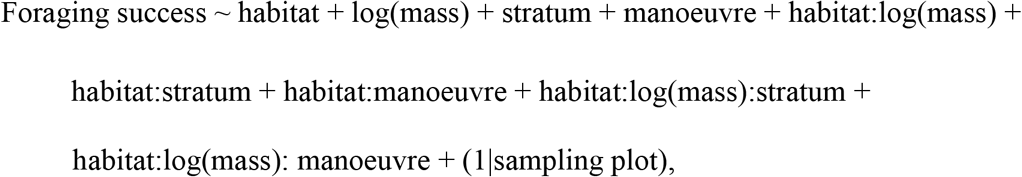

Where habitat was a factor variable with two levels (primary and logged), stratum was a factor variable with three levels (understorey, midstorey and canopy) and the manoeuvre was a factor variable with two levels (sally and non-sally; gleaners and hover-feeders were clubbed into a single category of birds feeding on prey in vegetation). Semicolons in the formulation represent interactions between predictor variables. The sampling plot was included as a random effect. All analyses were done in R (R Core Team, 2022).

## RESULTS

### Changes to the abiotic environment with selective logging

**Figure 2.**
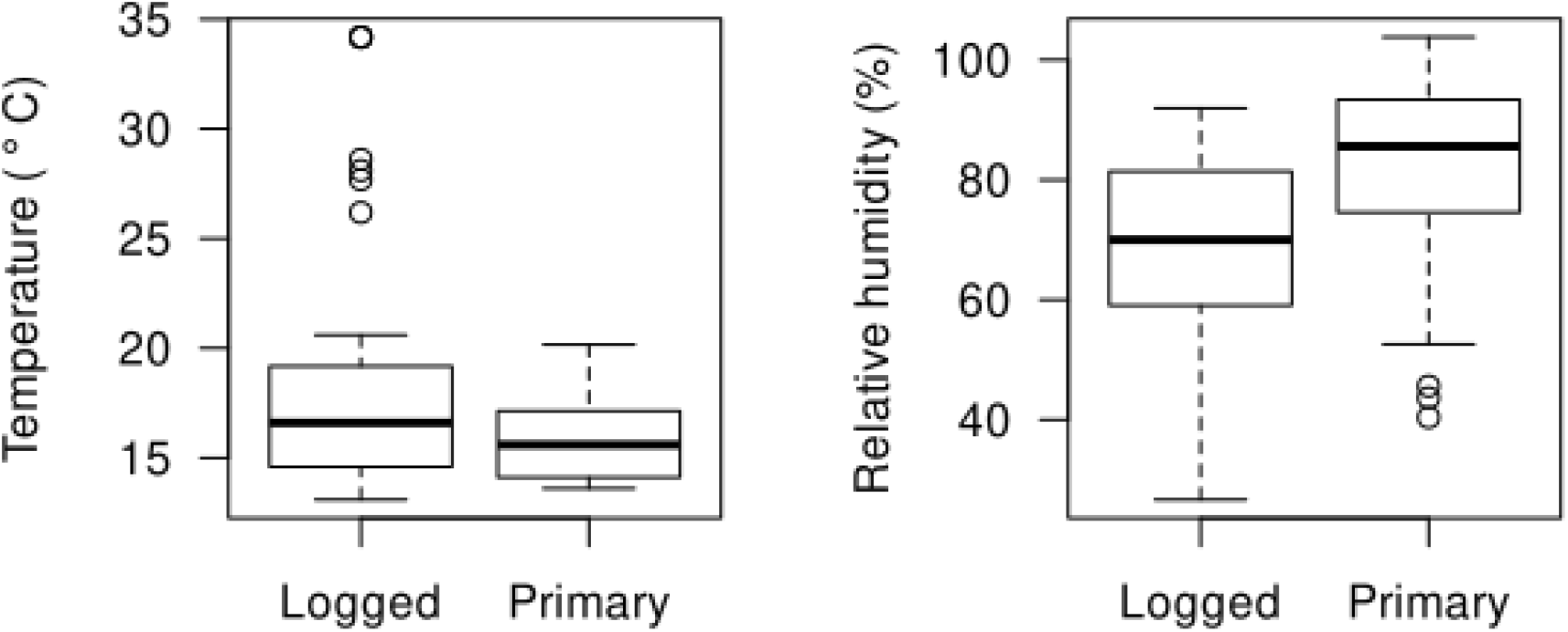
(a) Temperature (°C), and (b) relative humidity (%) in logged and primary forest. On average, primary forest was 2.3°C cooler and 14.6% more humid than logged forest (temperature: *F*_1,75_ = 4.43, p = 0.04, Fig. 2a; humidity: F_1,908_ = 282.3, p < 0.01, Fig. 2b).

### Selective logging and arthropod densities

We captured a total of 7,222 arthropods from the three different traps (primary forest: 4,198; logged forest: 3,024). Foliage arthropod density was higher in primary forest (branch beats; ANOVA, *F*_1,106_ = 4.22; *p* < 0.01; Fig.3) while density of arthropods in flight was higher in logged forest (sticky traps; Poisson ANOVA, *z*_1,106_ = -5.15; *p* < 0.01; Fig.3). There was no difference in the density of terrestrial arthropods in primary and logged forest (pitfall traps; Poisson ANOVA, *z*_1,106_ = 0.33; *p* = 0.75; Fig 3).

**Figure 3.**
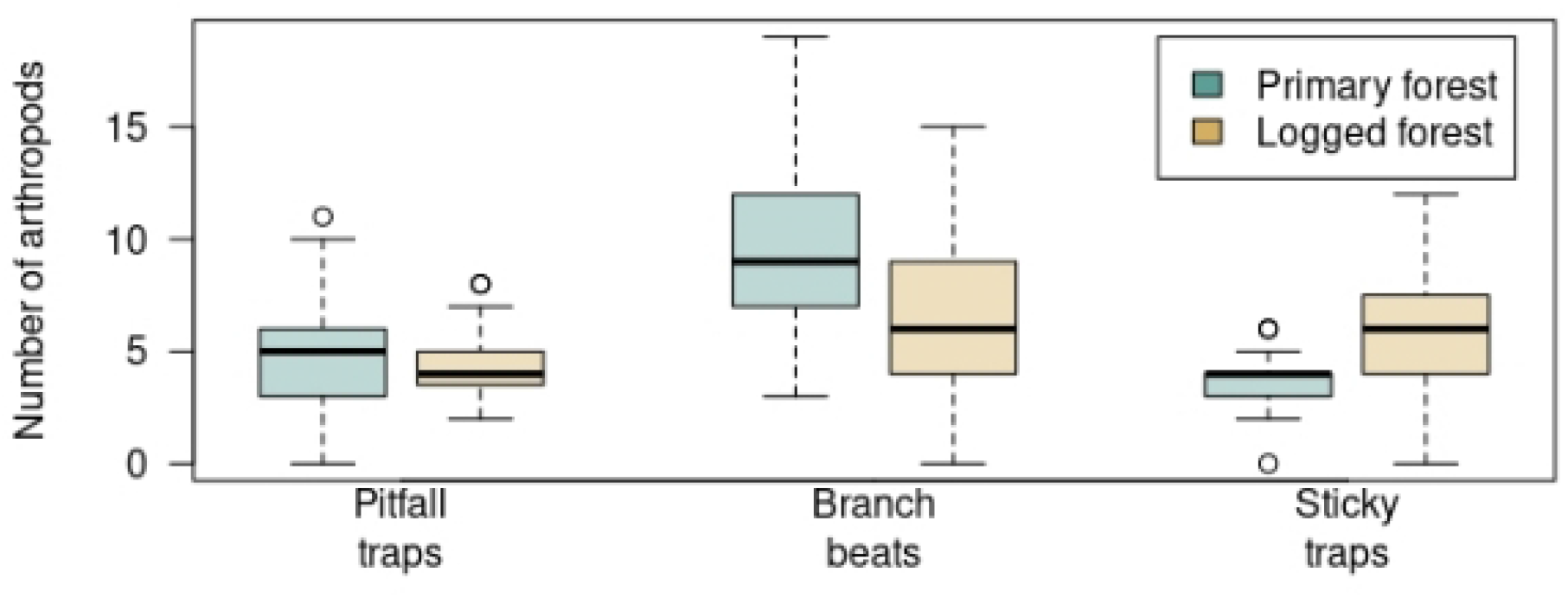
Arthropod densities in pitfall traps (terrestrial arthropods), branch beats (foliage arthropods) and sticky traps (flying arthropods) primary and logged forest.

### Arthropod community composition

We found that arthropod communities are largely composed differently in primary and logged forest Primary forest harbored more arthropods from the orders Arachnida (spiders, etc.) and Hemiptera (bugs), whereas logged forest had higher densities of Dipterans (flies; Fig. 4).

**Figure 4.**
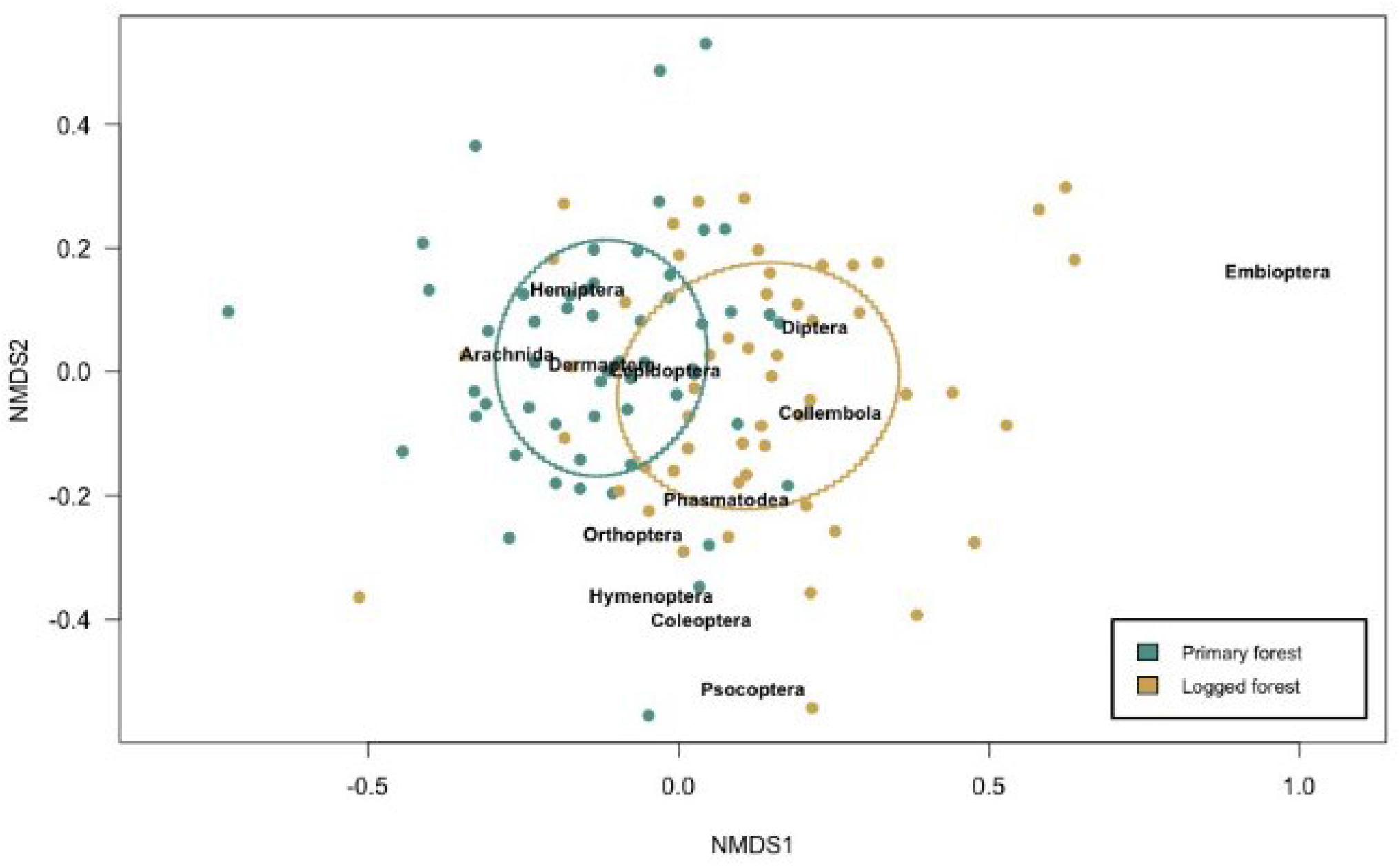
Non-metric multidimensional scaling (NMDS) ordination of arthropod orders shows differences in arthropod community composition between primary and logged forest (stress = 0.21)

### Foraging by birds in primary and logged forest

We observed 2,106 individual birds from 97 species for varying lengths of time (01 to 245 seconds, mean observation length = 38 seconds). Of these, 1,005 observations (with 2,065 instances of feeding attempts) were in primary forest and 1,101 (2,242 feeding attempts) were in logged forest. On limiting the data to only those species for which we had at least 10 observations in both primary and logged forest, we obtained data on 1,050 observations of 49 species, of which 593 observations were of birds attempting to feed, and 457 were non-feeding observations (e.g., birds preening or singing).

### Proportion of Time Spent Foraging by Birds in Primary and Logged Forest

Contrary to our expectations, we found that insectivorous birds spent roughly 10% more of their time foraging in logged than in primary forest (McFadden’s Pseudo-*R*_*2*_ from a quasibinomial generalised linear model = 0.44, Fig. 5). The higher time spent foraging in logged forest was independent of species traits such as body mass, foraging manoeuvre (gleaning or sallying) and foraging stratum (understory, midstorey or canopy).

**Figure 5.**
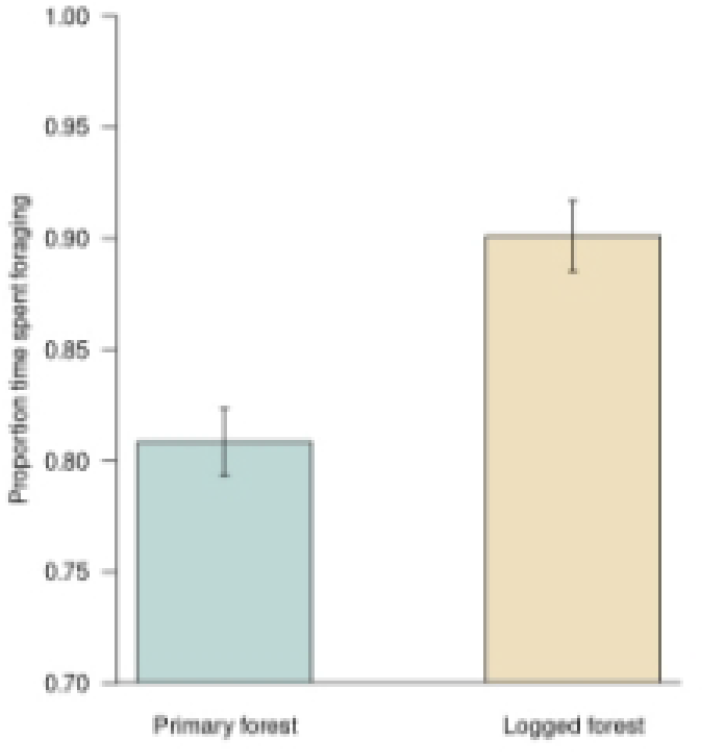
Proportion of time spent foraging by birds in primary and selectively logged forest.

### Foraging Success

Habitat (primary or logged), body mass, foraging manoeuvre and foraging stratum were all important determinants of foraging success (i.e., the proportion of attacks on prey that were successful; binomial generalised mixed-effects model with the plot as a random effect; marginal R_2_ = 0.26, conditional R_2_ = 0.30; Type II Analysis of Deviance table in Table S1; Fig. 6).

**Figure 6.**
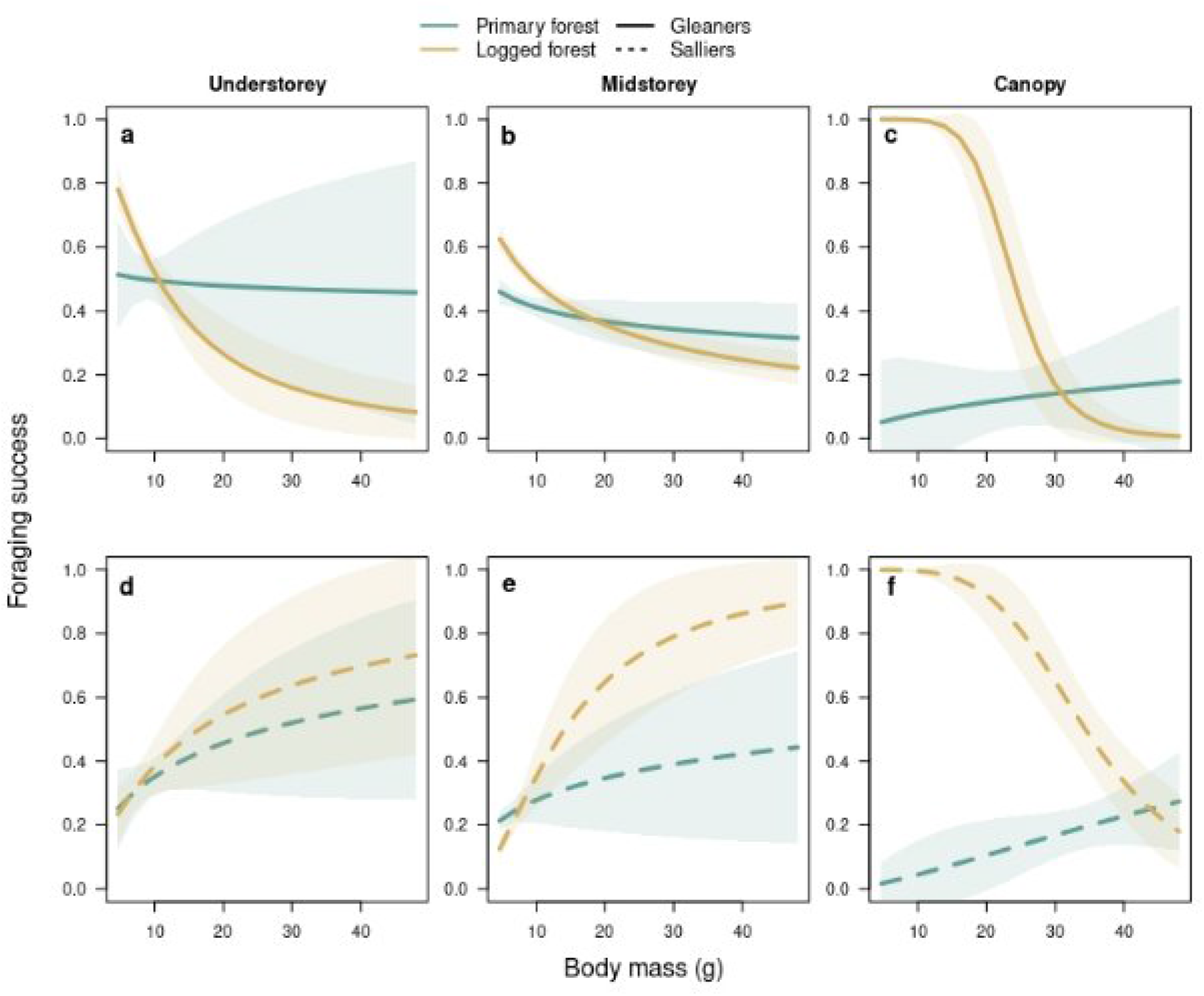
The relationship between foraging success and body mass (g) of insectivorous gleaners (continuous lines) and salliers (dotted lines) in primary (green) and logged (brown) forest in the understorey level (a, d); midstorey (b, e) and canopy (c, f).

For all understorey, midstorey and canopy gleaners foraging success did not vary with body mass in primary forest but declined sharply for larger species in the understorey and canopy of the logged forest and weakly in the midstorey of logged forest (Fig. 6a). Salliers showed stronger patterns of foraging success with body size in logged than in primary forest (Fig. 6d-f). In both the understorey and midstorey, larger salliers had greater foraging success than smaller salliers (Fig. 6d, e), a pattern opposite to that observed for gleaners (Fig. 6a, b).

## DISCUSSION

Our results indicate that selective logging has important impacts on temperature-humidity microclimates, the abundances of different types of arthropods and on both the time spent foraging and foraging success of insectivorous bird species. Insectivorous birds are particularly threatened by all forms of forest degradation (Powell et al., 2015). Here, we show that foraging success differs variably with body masses of insectivorous gleaners and salliers at different strata (understorey, midstorey and canopy) of logged and primary forest depending on the availability of prey arthropods.

Surprisingly, we found that birds spent more time foraging in logged than in primary forest, contrary to our prediction that warmer temperatures in logged forest would limit foraging activity because of thermal stress. While warmer temperatures in the logged forest might potentially lead to thermal stress, increasing temperatures and the decline in humidity with logging might concurrently result in a lower abundance of arthropod food resources, which we report here for foliage arthropods (Fig. 4). The diets of foliage gleaners are dominated by lepidopterans (butterflies and moths), hemipterans (bugs) and arachnids (especially spiders) (Supriya et al., 2020). We show a reduced abundances of arthropods from these orders with logging. Most birds in logged forest might therefore have to spend more time and energy foraging in order to meet food requirements compared with birds in primary forest. Spending a greater amount of time foraging is likely to come at the cost of time allocated to other activities such as reproductive effort. The necessity to expend greater energy in obtaining sufficient food might underlie observed patterns in body condition (for almost all species, individuals are significantly lighter in logged than in primary forest; Srinivasan & Wilcove, 2021), and consequently demographic performance in terms of survival and recruitment (Srinivasan, 2019; Srinivasan & Wilcove, 2021).

Temperature and humidity play an important role in the regulation of arthropod diversity and biomass (Savopoulou-Soultani et al., 2012). Being ectotherms, arthropods are highly dependent on external temperature and moisture for normal physiology (Bale et al., 2002; Menéndez, 2007; Jaworski & Hilszczański., 2013). Invertebrates also tend to have narrow thermal and moisture tolerances (Deutsch et al., 2008, Højer et al. 2001) and are likely to deal poorly with changes in temperature and humidity (Peck et al., 2008) and.). Changes in these abiotic variables with logging could therefore impact arthropod availability for insectivorous bird species. We found that the density of arthropods in foliage (fed on by gleaning birds) was far higher in primary than in logged forest but found the opposite pattern for flying insects (fed on by sallying birds), which were significantly more common in logged forest. These patterns are limited to only the understorey and midstorey strata of the forest; we did not sample canopy arthropods.

The patterns in understorey-midstorey arthropod abundance are consistent with the finding that, in general, understorey and midstorey gleaners have greater foraging success in primary forest where foliage arthropods are much more common while understorey salliers are more successful in catching prey in logged forest, where arthropods in flight are more abundant. Given that arthropod abundance is likely to drive foraging behaviour and success, the patterns in foraging success we report for canopy species might also result from changes in canopy structure, microclimates and arthropod availability. Logging can have a negative impact on canopy arthropod abundance (Turner and Foster, 2009). The loss of vegetation in the canopy of logged forest reduces foliage volume in the canopy, and therefore also should reduce foraging success for canopy gleaners, especially large canopy gleaners with greater resource requirements (Fig. 6c). Small salliers in the canopy of the logged forest had higher foraging success than small canopy salliers in primary forest (Fig. 6f). However, foraging success for canopy salliers declined drastically with increasing body size in logged forest, while foraging success of canopy salliers increased with increasing body mass in primary forest (Fig. 6f).

Selective logging might increase the abundance of small flying insects in general and result in a reduction in the density of large flying insects. Such a change in the pattern of availability of insect prey might explain why – in the canopy – smaller salliers have high foraging success in logged forest, but larger salliers have higher foraging success in primary forest (Fig. 6f). Future work should test whether there is a size-dependent change in the density of flying insects with logging, potentially explaining the patterns in foraging success that we report.

A further possibility is that different changes in vegetation structure and composition in different vertical strata in the logged forest – e.g., increased volume of understory, variable effects in the midstorey and thinning of the canopy – can also have variable effects on the abundance, composition and diversity of arthropod prey of various types –large versus small, prey in foliage versus prey in flight, etc. Open and sunny patches in the logged forest might increase habitat use by flying insects, thereby leading to more foraging success by understorey and midstorey salliers.

Our work examines the foraging behaviour of montane birds in the eastern Himalayas with the aim of understanding the role of a pervasive form of land-use change on one of the important drivers of individual fitness of birds.

## ACKNOWLEDGEMENTS

We thank the Arunachal Pradesh Forest Department and the Shergaon Forest Division for their continued support of this project and for providing us with permits to conduct this work. We thank Shreesh Kaulgud, Paulami Sarkar and Supriya Samanta for their help with data collection and processing.

## Ethics statement

We thank the Forest Department of the state of Arunachal Pradesh for providing permissions for this study (permit no. CWL/G/173/2018-2019/Pt-VII(A)/47-48). The Institutional Animal Ethics Committee of the Indian Institute of Science approved this study.

## Author contributions

KA and US co-conceived the study, KA, RC, SR, MR, DKP, DS, AB and BT collected the data. KA, RC and US analysed the data. KA wrote the manuscript, and all authors contributed to its revision.

## Data depository

Data will be archived on Data Dryad upon acceptance.

## Supplementary Material

**Table S1.** Type II Analysis of Deviance Table for predictors used to model foraging success using a binomial generalised linear mixed-effects model.

| Predictor variable | Likelihood<br>Ratio $\chi^2$ | Degrees of freedom | | P |
| --- | --- | --- | --- | --- |
| Habitat (primary or logged) | 3.06 | 1 |  | 0.08 |
| Log (body mass) | 13.28 | 1 |  | < 0.01 |
| Stratum (understorey, midstorey or canopy) | 9.33 | 2 |  | < 0.01 |
| Foraging manoeuvre (glean or sally) | 99.97 | 1 |  | < 0.01 |
| Habitat-Log (body mass) interaction | 5.65 | 1 |  | 0.02 |
| Habitat-Stratum interaction | 5.19 | 2 |  | 0.07 |
| Habitat-Foraging manoeuvre interaction | 9.55 | 1 |  | < 0.01 |
| Habitat-Log (body mass)-Stratum interaction | 7.53 | 4 | 0.11 |  |
| Habitat-Log (body mass)-Manoeuvre interaction | 13.88 | 2 | <0.01 |  |

## LITERATURE CITED

Bale, J. S., Masters, G. J., Hodkinson, I. D., Awmack, C., Bezemer, T. M., Brown, V. K., Butterfield, J., Buse, A., Coulson, J. C., & Farrar, J. (2002). Herbivory in global climate change research: direct effects of rising temperature on insect herbivores. Global Change Biology, 8(1), 1–16.

Basset, Y., Cizek, L., Cuénoud, P., Didham, R. K., Guilhaumon, F., Missa, O., Novotny, V., Ødegaard, F., Roslin, T., & Schmidl, J. (2012). Arthropod diversity in a tropical forest. Science, 338(6113), 1481–1484.

Basset, Y., Charles, E., Hammond, D. S., & Brown, V. K. (2001). Short-term effects of canopy openness on insect herbivores in a rain forest in Guyana. Journal of Applied Ecology, 38(5), 1045–1058.

Bregman, T. P., Sekercioglu, C. H., & Tobias, J. A. (2014). Global patterns and predictors of bird species responses to forest fragmentation: implications for ecosystem function and conservation. Biological Conservation, 169, 372–383.

Burivalova, Z., Lee, T. M., Giam, X., Şekercioğlu, Ç. H., Wilcove, D. S., & Koh, L. P. (2015). Avian responses to selective logging shaped by species traits and logging practices. Proceedings of the Royal Society B: Biological Sciences, 282(1808), 20150164.

Canaday, C., & Rivadeneyra, J. (2001). Initial effects of a petroleum operation on Amazonian birds: terrestrial insectivores retreat. Biodiversity & Conservation, 10(4), 567–595.

Chung, A. Y. C., Eggleton, P., Speight, M. R., Hammond, P. M., & Chey, V. K. (2000). The diversity of beetle assemblages in different habitat types in Sabah, Malaysia. Bulletin of entomological research, 90(6), 475–496.

Cornelius, M. L., & Osbrink, W. L. (2010). Effect of soil type and moisture availability on the foraging behavior of the Formosan subterranean termite (Isoptera: Rhinotermitidae). Journal of Economic Entomology, 103(3), 799–807.

Davis, A. J. (2000). Does reduced-impact logging help preserve biodiversity in tropical rainforests? A case study from Borneo using dung beetles (Coleoptera: Scarabaeoidea) as indicators. Environmental Entomology, 29(3), 467–475.

Deutsch, C. A., Tewksbury, J. J., Huey, R. B., Sheldon, K. S., Ghalambor, C. K., Haak, D. C., & Martin, P. R. (2008). Impacts of climate warming on terrestrial ectotherms across latitude. Proceedings of the National Academy of Sciences, 105(18), 6668–6672.

Edwards, D. P., & Laurance, W. F. (2013). Biodiversity despite selective logging. Science, 339(6120), 646–647.

Elsen, P. R., Monahan, W. B., & Merenlender, A. M. (2018). Global patterns of protection of elevational gradients in mountain ranges. Proceedings of the National Academy of Sciences, 115(23), 6004–6009.

Ewers, R. M., Boyle, M. J., Gleave, R. A., Plowman, N. S., Benedick, S., Bernard, H., Bishop, T. R., Bakhtiar, E. Y., Chey, V. K., & Chung, A. Y. (2015). Logging cuts the functional importance of invertebrates in tropical rainforest. Nature Communications, 6(1), 1–7.

FAO and UNEP. 2020. (2020). The State of the World’s Forests 2020 (ISBN: 978-92-5-132419-6). Forests, biodiversity and people. Rome.

Freeman, B. G., Song, Y., Feeley, K. J., & Zhu, K. (2021). Montane species track rising temperatures better in the tropics than in the temperate zone. Ecology Letters, 24(8), 1697–1708.

Grenyer, R., Orme, C. D. L., Jackson, S. F., Thomas, G. H., Davies, R. G., Davies, T. J., Jones, K. E., Olson, V. A., Ridgely, R. S., & Rasmussen, P. C. (2006). Global distribution and conservation of rare and threatened vertebrates. Nature, 444(7115), 93–96.

Hill, J. K., Hamer, K. C., Lace, L. A., & Banham, W. M. T. (1995). Effects of selective logging on tropical forest butterflies on Buru, Indonesia. Journal of Applied Ecology, 754–760.

Hill, J. K. (1999). Butterfly spatial distribution and habitat requirements in a tropical forest: impacts of selective logging. Journal of Applied Ecology, 36(4), 564–572.

Højer, R., Bayley, M., And, C. F. D., & Holmstrup, M. (2001). Stress synergy between drought and a common environmental contaminant: studies with the collembolan Folsomia candida. Global Change Biology, 7(4), 485–494.

Holloway, J. D., Kirk-Spriggs, A. H., & Khen, C. V. (1992). The response of some rain forest insect groups to logging and conversion to plantation. Philosophical Transactions of the Royal Society of London. Series B: Biological Sciences, 335(1275), 425–436.

Hamilton, A. J., Basset, Y., Benke, K. K., Grimbacher, P. S., Miller, S. E., Novotnỳ, V., Samuelson, G. A., Stork, N. E., Weiblen, G. D., & Yen, J. D. (2010). Quantifying uncertainty in estimation of tropical arthropod species richness. The American Naturalist, 176(1), 90–95.

Jaworski, T., & Hilszczański, J. (2013). The effect of temperature and humidity changes on insects development their impact on forest ecosystems in the context of expected climate change.

Menéndez, R. (2007). How are insects responding to global warming? Tijdschrift Voor Entomologie, 150(2), 355.

Oksanen, J., Blanchet, F. G., Friendly, M., Kindt, R., Legendre, P., McGlinn, D., Minchin, P. R., O’Hara, R. B., Simpson, G. L., Solymos, P., Stevens, M. H. H., Szoecs, E., & Wagner, H. (2019). vegan: Community Ecology Package. R Foundation for Statistical Computing.

Pavlacky Jr, D. C., Possingham, H. P., & Goldizen, A. W. (2015). Integrating life history traits and forest structure to evaluate the vulnerability of rainforest birds along gradients of deforestation and fragmentation in eastern Australia. Biological Conservation, 188, 89–99.

Peck, L. S., Webb, K. E., Miller, A., Clark, M. S., & Hill, T. (2008). Temperature limits to activity, feeding and metabolism in the Antarctic starfish Odontaster validus. Marine Ecology Progress Series, 358, 181–189.

Pimm, S. L. (2008). Biodiversity: climate change or habitat loss—which will kill more species? Current Biology, 18(3), R117–R119.

Powell, L. L., Cordeiro, N. J., & Stratford, J. A. (2015). Ecology and conservation of avian insectivores of the rainforest understory: A pantropical perspective. Biological Conservation, 188, 1–10.

Pandit, M. K., Sodhi, N. S., Koh, L. P., Bhaskar, A., & Brook, B. W. (2007). Unreported yet massive deforestation driving loss of endemic biodiversity in Indian Himalaya. Biodiversity and Conservation, 16(1), 153–163.

Peh, K. S.-H., de Jong, J., Sodhi, N. S., Lim, S. L.-H., & Yap, C. A.-M. (2005). Lowland rainforest avifauna and human disturbance: persistence of primary forest birds in selectively logged forests and mixed-rural habitats of southern Peninsular Malaysia. Biological Conservation, 123(4), 489–505.

Powell, L. L., Stouffer, P. C., & Johnson, E. I. (2013). Recovery of understory bird movement across the interface of primary and secondary Amazon rainforest. The Auk, 130(3), 459–468.

R Core Team (2021). R: A language and environment for statistical computing. R Foundation for Statistical Computing, Vienna, Austria. URL https://www.R-project.org/.

Robinson, S. K., & Holmes, R. T. (1982). Foraging behavior of forest birds: the relationships among search tactics, diet, and habitat structure. Ecology, 63(6), 1918–1931.

Rutt, C. L., Jirinec, V., Cohn-Haft, M., Laurance, W. F., & Stouffer, P. C. (2019). Avian ecological succession in the Amazon: A long-term case study following experimental deforestation. Ecology and Evolution, 9(24), 13850–13861.

Savopoulou-Soultani, M., Papadopoulos, N. T., Milonas, P., & Moyal, P. (2012). Abiotic factors and insect abundance. Hindawi.

Senior, R. A., Hill, J. K., González del Pliego, P., Goode, L. K., & Edwards, D. P. (2017). A pantropical analysis of the impacts of forest degradation and conversion on local temperature. Ecology and Evolution, 7(19), 7897–7908.

Senior, R. A., Hill, J. K., Benedick, S., & Edwards, D. P. (2018). Tropical forests are thermally buffered despite intensive selective logging. Global Change Biology, 24(3), 1267–1278.

Srinivasan, U., Hines, J. E., & Quader, S. (2015). Demographic superiority with increased logging in tropical understorey insectivorous birds. Journal of Applied Ecology, 52(5), 1374–1380.

Srinivasan, U. (2019). Morphological and behavioral correlates of long-term bird survival in selectively logged forest. Frontiers in Ecology and Evolution, 17.

Srinivasan, U., & Wilcove, D. S. (2021). Interactive impacts of climate change and land-use change on the demography of montane birds. Ecology, 102(1), e03223.

Stratford, J. A., & Stouffer, P. C. (1999). Local extinctions of terrestrial insectivorous birds in a fragmented landscape near Manaus, Brazil. Conservation Biology, 13(6), 1416–1423.

Stratford, J. A., & Robinson, W. D. (2005). Gulliver travels to the fragmented tropics: geographic variation in mechanisms of avian extinction. Frontiers in Ecology and the Environment, 3(2), 85–92.

Stouffer, P. C., Jirinec, V., Rutt, C. L., Bierregaard Jr, R. O., Hernández-Palma, A., Johnson, E. I., Midway, S. R., Powell, L. L., Wolfe, J. D., & Lovejoy, T. E. (2021). Long-term change in the avifauna of undisturbed Amazonian rainforest: ground-foraging birds disappear and the baseline shifts. Ecology Letters, 24(2), 186–195.

Supriya, K., Price, T. D., & Moreau, C. S. (2020). Competition with insectivorous ants as a contributor to low songbird diversity at low elevations in the eastern Himalaya. Ecology and Evolution, 10(10), 4280–4290.

Turner, E. C., & Foster, W. A. (2009). The impact of forest conversion to oil palm on arthropod abundance and biomass in Sabah, Malaysia. Journal of Tropical Ecology, 25(1), 23–30.

Vasconcelos, H. L., Vilhena, J. M. S., & Caliri, G. J. A. (2000). Responses of ants to selective logging of a central Amazonian forest. Journal of Applied Ecology, 37(3), 508–514.

Willett, T. R. (2001). Spiders and other arthropods as indicators in old-growth versus logged redwood stands. Restoration Ecology, 9(4), 410–420.

Willott, S. J., Lim, D. C., Compton, S. G., & Sutton, S. L. (2000). Effects of selective logging on the butterflies of a Bornean rainforest. Conservation Biology, 14(4), 1055–1065.

